# Unravelling the hidden diversity of the monkfish *Lophius gastrophysus* using integrative taxonomy

**DOI:** 10.64898/2026.11.10.765400

**Authors:** Ananda Krishna Pereira D’Elia, Leticia Maria Monteiro Soares, Frederico Henning, Antonio Mateo Solé-Cava

**Affiliations:** Centro Nacional para a Identificação Molecular do Pescado (CENIMP), Departamento de Genética, Instituto de Biologia, Universidade Federal do Rio de Janeiro (UFRJ), Rio de Janeiro, RJ, Brazil

**Keywords:** Anglerfish, Automatic Species Delimitation Methods, Cryptic species, Fisheries, Geometric Morphometry, Lophiidae

## Abstract

The monkfish *Lophius gastrophysus* is a commercially exploited demersal species distributed along the continental shelf and slope of the western Atlantic Ocean. Like other anglerfish species, *L. gastrophysus* has a pelagic larval stage that can facilitate dispersal by oceanic currents (Fariña et al., 2008). It is considered the most widely distributed species in the genus, spanning more than 10,000 km of the western Atlantic Coast, from North Carolina to Argentina. Previous morphological analyses based on a limited set of traits suggested that *L. gastrophysus* may comprise two distinct species. We tested this hypothesis by combining molecular and morphometric data. Sequence data were obtained from 38 *L. gastrophysus* individuals collected from Northern and Southern American populations. We also used a subset of 13 specimens for geometric morphometrics. Phylogenetic analyses and automatic species delimitation methods consistently distinguished Northern and Southern Atlantic *L. gastrophysus* as separate clades, and geometric morphometrics supported the same split. The congruence among geographic, molecular, and morphometric data indicates that *L. gastrophysus* harbours two distinct lineages, highlighting the need for a formal taxonomic revision and separate fisheries management of northern and southern Atlantic populations.

## 1 INTRODUCTION

The West Atlantic monkfish, *Lophius gastrophysus* (Miranda Ribeiro, 1915), inhabits the continental shelf and slope of the western Atlantic Ocean, reportedly between North Carolina and Argentina (Caruso, 1983), with recent records expanding its known distribution into the south-western Gulf of Mexico (Del Moral-Flores et al., 2024). This makes it the most widely distributed species in the genus (Grant & Leslie, 1993). The species became a successful target for commercial exploitation in 2000 (Valentim et al., 2007), escalating so rapidly that, by 2001, the removal rate of monkfish had reached 16% of the total stock (Perez, 2005). In response to declining catch rates, fishing pressure was intensified to maintain yields, ultimately threatening stock sustainability through overexploitation. This scenario led to the establishment of legal measures to regulate the targeted fishery for this species in Brazil (Valentim et al., 2007).

Consistent morphometric differences, such as the length of the illicium and the size of the pair of dark tendrils on the snout, have been reported between *Lophius gastrophysus* populations from the North and South Atlantic (Caruso 1983). However, Caruso (1983) cautioned that relying solely on these two morphological characteristics could not provide sufficient evidence to identify two different species, given inadequate sampling of the Southern population. The advent of molecular data has allowed the delimitation of species that are otherwise morphologically indistinguishable (Blaxter, 2004; Tautz et al., 2002). Conflating cryptic species because identification resolution is insufficient leads to inaccurate management policies, which can be especially threatening to economically relevant species (Longo et al., 2022; Fišer et al., 2018). Integrative taxonomic approaches have proven highly effective in resolving these taxonomic uncertainties, uncovering unanticipated hidden diversity even within closely related goosefish genera such as *Lophiomus* (Chen et al., 2024). Cryptic species are commonly attributed to recent divergence, morphological convergence or parallelism through phylogenetic niche conservatism (Fišer et al., 2018). The latter mechanism is often associated with strong stabilising selection under extreme environmental conditions, which limits morphological variation among species (Janssen et al., 2015). Because deep-sea habitats typically have challenging conditions (e.g., low oxygen concentrations, high pressure, and low luminosity), these environments may harbour a rich diversity of cryptic species (Bickford et al., 2007).

Given the economic importance of West Atlantic monkfish and the serious consequences of inaccurate delimitation of its already depleted stocks (Silva et al., 2022), assessing genetic and morphological variation across its distribution is imperative. Here, we applied an integrative species delimitation approach – combining phylogenetic analyses of nuclear and mitochondrial sequences, automatic species delimitation methods (SDMs) and geometric morphometrics – to demonstrate that *Lophius cf. gastrophysus* harbours two very distinct lineages, one in the northern and one in the southern West Atlantic.

## 2 MATERIAL AND METHODS

### 2.1 Sampling

We analysed 38 muscle or fin tissue samples of *Lophius gastrophysus* (Table 1), including 29 from South America and nine from North America (Figure 1). As outgroups, we also included three samples of *L. americanus* (Valenciennes, 1837) from the USA, four of the European *L. budegassa* (Spinola, 1807), and one of the European *L. piscatorius* (Linnaeus, 1758). For fresh samples, approximately 50 mg of muscle or fin clip tissue from each fish was collected, preserved in absolute ethanol, and stored at -20 °C for downstream analyses. North American tissue samples were obtained as a loan from the Ichthyology Collection of the Biodiversity Institute & Natural History Museum of the University of Kansas (KU).

**Figure 1.**
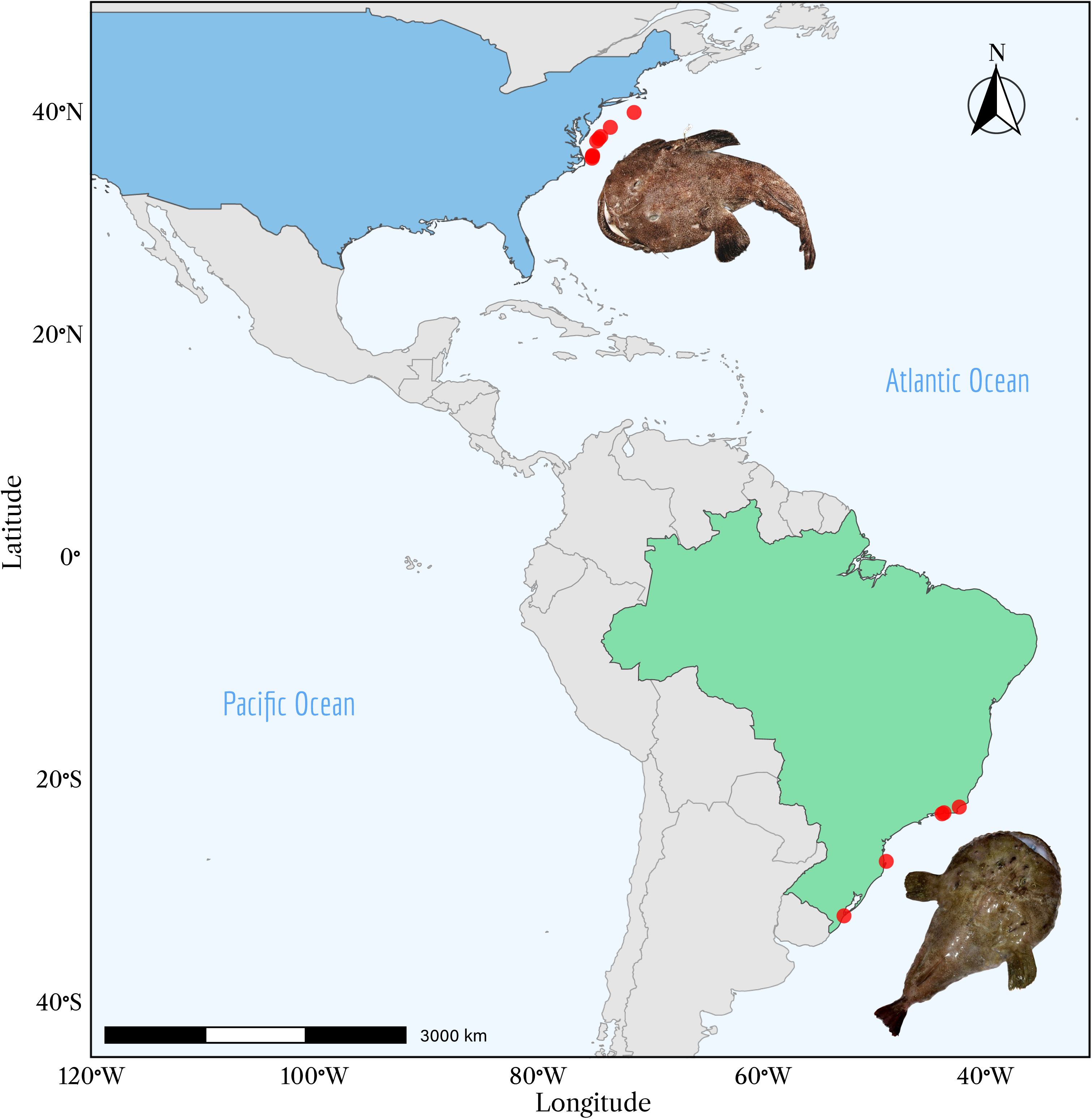
Sampling locations in the USA (blue) and Brazil (green) from which *Lophius cf. gastrophysus* samples were obtained. Photo of the MCZ 159679 specimen was kindly provided by the Museum of Comparative Zoology (Harvard).

**Table 1.** Taxon sampling and corresponding locality and field/museum number for *Lophius gastrophysus* and outgroup species. Specimens marked with a symbol (†) were used in the geometric morphometric analyses. For the museum samples, catalogue numbers are as follows: KU 4956 to KU 4961= MCZ159678 to MCZ159683 respectively; KU 8203 = MCZ166186 and KU 8204 = MCZ166187.

| **Species** | **Locality** | **Field number** |
|---|---|---|
| *L. gastrophysus* | Glória, Rio de Janeiro, RJ, Brazil | AD01 |
| *L. gastrophysus* | Barra da Tijuca, Rio de Janeiro, RJ, Brazil | AD02† |
| *L. gastrophysus* | Barra da Tijuca, Rio de Janeiro, RJ, Brazil | AD03† |
| *L. gastrophysus* | Barra da Tijuca, Rio de Janeiro, RJ, Brazil | AD04† |
| *L. gastrophysus* | Barra da Tijuca, Rio de Janeiro, RJ, Brazil | AD05† |
| *L. gastrophysus* | Barra da Tijuca, Rio de Janeiro, RJ, Brazil | AD06† |
| *L. gastrophysus* | Macaé, Rio de Janeiro, RJ, Brazil | AT23 |
| *L. gastrophysus* | Macaé, Rio de Janeiro, RJ, Brazil | AT24 |
| *L. gastrophysus* | Macaé, Rio de Janeiro, RJ, Brazil | AT25 |
| *L. gastrophysus* | Macaé, Rio de Janeiro, RJ, Brazil | AT26 |
| *L. gastrophysus* | Macaé, Rio de Janeiro, RJ, Brazil | AT27 |
| *L. gastrophysus* | Macaé, Rio de Janeiro, RJ, Brazil | AT28 |
| *L. gastrophysus* | Macaé, Rio de Janeiro, RJ, Brazil | AT29 |
| *L. gastrophysus* | Macaé, Rio de Janeiro, RJ, Brazil | AT30 |
| *L. gastrophysus* | Macaé, Rio de Janeiro, RJ, Brazil | AT31 |
| *L. gastrophysus* | Macaé, Rio de Janeiro, RJ, Brazil | AT32 |
| *L. gastrophysus* | São Sebastião-Arvoredo, Santa Catarina, SC, Brazil | E61 |
| *L. gastrophysus* | São Sebastião-Arvoredo, Santa Catarina, SC, Brazil | E62 |
| *L. gastrophysus* | São Sebastião-Arvoredo, Santa Catarina, SC, Brazil | E63 |
| *L. gastrophysus* | São Sebastião-Arvoredo, Santa Catarina, SC, Brazil | E64 |
| *L. gastrophysus* | São Sebastião-Arvoredo, Santa Catarina, SC, Brazil | E65 |
| *L. gastrophysus* | São Sebastião-Arvoredo, Santa Catarina, SC, Brazil | E66 |
| *L. gastrophysus* | São Sebastião-Arvoredo, Santa Catarina, SC, Brazil | E67 |
| *L. gastrophysus* | São Sebastião-Arvoredo, Santa Catarina, SC, Brazil | E68 |
| *L. gastrophysus* | Rio Grande do Sul, RS, Brazil | RS1 |
| *L. gastrophysus* | Rio Grande do Sul, RS, Brazil | RS2 |
| *L. gastrophysus* | Rio Grande do Sul, RS, Brazil | RS3 |
| *L. gastrophysus* | Rio Grande do Sul, RS, Brazil | RS4 |
| *L. gastrophysus* | Rio Grande do Sul, RS, Brazil | RS5 |
| *L. gastrophysus* | Slope, Mid Atlantic Bight, 36.0666700000°, -74.8000000000°, USA | KU 3265 |
| *L. gastrophysus* | Slope, south New England, 36.1669444000°, -74.7741667000°, USA | KU 4956† |
| *L. gastrophysus* | Slope, south New England, 37,6727778000°, -74,2097222000°, USA | KU 4957† |
| *L. gastrophysus* | Slope, south New England, 38,6883333000°, -73,1766667000°, USA | KU 4958† |
| *L. gastrophysus* | Slope, south New England, 35,9116667000°, -74,8216667000°, USA | KU 4959† |
| *L. gastrophysus* | Slope, south New England, 37.8588889000°, -74.0430556000°, USA | KU 4960† |
| *L. gastrophysus* | Slope, south New England, 36.0716667000°, -74.7766667000°, USA | KU 4961† |
| *L. gastrophysus* | Slope, 37,4500000000°, -74,4083333000°, USA | KU 8203† |
| *L. gastrophysus* | Mid Atlantic Bight, 40.0216667000°, -71.0416667000°, USA | KU 8204† |
| *L. budegassa* | Golfe de Fos, Provence-Alpes-Côte d'Azur, France | BUD1 |
| *L. budegassa* | Golfe de Fos, Provence-Alpes-Côte d'Azur, France | BUD2 |
| *L. budegassa* | Golfe de Fos, Provence-Alpes-Côte d'Azur, France | BUD3 |
| *L. budegassa* | Golfe de Fos, Provence-Alpes-Côte d'Azur, France | BUD4 |
| *L. americanus* | Manhattan, New York, United States | AME1 |
| *L. americanus* | Manhattan, New York, United States | AME2 |
| *L. americanus* | Manhattan, New York, United States | AME3 |
| *L. piscatorius* | Konstanz, Germany | PISC1 FH |

### 2.2 Molecular analyses

#### DNA extraction, amplification, and sequencing

Genomic DNA was extracted using the salting-out method described by Miller et al. (1988). Two mitochondrial genes, cytochrome C oxidase 1 (*COI*) and cytochrome b (*Cytb*), and one nuclear gene, rhodopsin (*Rod*), were amplified by PCR. Primer sets were Fish-F1 and Fish-R1(Ward et al., 2005) for *COI*, GluFish-F and THR-Fish2-R (Sevilla et al., 2007) for *Cytb* and Rod-F2W and Rod-R4n (Sevilla et al., 2007) for *Rod*. The cycling conditions consisted of an initial denaturation at 95°C for 4 min, followed by 33 cycles of denaturation at 95°C for 30 s, annealing at 56°C (*COI*) or 55°C (*Cytb* and *Rod*) for 1 min, extension at 72°C for 40 s, and a final extension at 72°C for 4 min. PCR reactions were conducted in a total reaction volume of 15 μL containing 0.5 μM of each primer, 1U cTaq DNA Polymerase Hot Start (Cellco), 200 μM dNTPs, and 5-10 ng of DNA template in Taq HS Reaction Buffer. PCR products were purified by adding 0.7 units of shrimp alkaline phosphatase (rSAP, Cellco) and 7 units of Exonuclease 1 (Exo1, Cellco) followed by incubation at 37 °C for one hour, to remove dNTPs and primers. Purified PCR products were sequenced using the BigDye Terminator v3.1 Cycle Sequencing Kit (Thermo Fisher Scientific, Applied Biosystems) and a SeqStudio Sequence Analyzer (Thermo Fisher Scientific, Applied Biosystems).

#### Phylogenetic analysis and distance estimation

Trace files were manually curated in Geneious Prime® v. 2026.0.2 (Kearse et al., 2012). Final sequence matrices consisted of sequences obtained here, complemented by other sequences of *Lophius,* and the outgroups *Lophiomus* Gill, 1883 and *Lophiodes* Goode & Bean, 1896 available in GenBank (Table S1, Table S2, and Table S3, Supporting Information 1). The final sequence matrices of *COI, Cytb,* and Rod comprised 87, 93 and 64 sequences, respectively. Consensus sequences were aligned with the default parameters of MAFFT v7.475 (Katoh & Standley, 2013). Maximum-likelihood phylogenetic inferences for *COI*, *Cytb* and *Rod* were performed using IQ-TREE (Nguyen et al., 2015) with 1,000 ultrafast bootstrap replicates to assess branch support. The best-fit nucleotide substitution models were automatically selected using ModelFinder as implemented in IQ-TREE. For the *COI* and *Cytb* datasets, the TN+F+G4 model (Tamura & Nei, 1993) was selected. In contrast, for the rhodopsin (*Rod*) dataset, the HKY+F model (Hasegawa et al., 1985) was identified as the best-fit model according to the Bayesian Information Criterion (BIC).

Average intra- and interspecific distances for *Lophius* nominal species and between samples of *L. gastrophysus l.s.* from the North and South Atlantic were computed for each final sequence matrix using MEGA X (Kumar et al., 2018) under the TN93 substitution model, which was identified as the best-fit model according to the Bayesian Information Criterion (BIC).

#### Secondary species delimitation analyses

Automatic species delimitation was performed using three methods: ASAP (Assemble Species by Automatic Partitioning; Puillandre et al., 2021), mPTP (Multi-rate Poisson Tree Processes; Kapli et al., 2017) and GMYC (Generalized Mixed Yule Coalescent Model, Fujisawa and Barraclough, 2013). The ASAP analyses were conducted on the Spart Explorer web server (https://spartexplorer.mnhn.fr/), using default parameters for relative gap width (1.5), Pmin (0.001), and Pmax (0.1). We used 10 steps for each run, with distances based on the Kimura (1980) two-parameter substitution model. We performed multi-rate Poisson Tree Processes (mPTP) analyses on the mPTP web server (http://mptp.h-its.org), applying the maximum likelihood heuristic and the multi-rate coalescent model under default settings. We used a minimum branch length threshold of 0.0001 and an outgroup for tree rooting, then excluded it from the delimitation analysis using the --outgroup_crop option.

For GMYC runs, we first generated Bayesian ultrametric trees for each locus dataset (*COI*, *Cytb,* and *Rod*) using BEAUTI 10.5.0 (Bouckaert et al., 2014) with the Yule model as the tree prior, then performed Bayesian phylogenetic inference using BEAST 10.5.0 (Bouckaert et al., 2014). Convergence and Effective Sample Sizes (ESS) were confirmed using Tracer 1.7.2 (Rambaut et al., 2018). The consensus trees were annotated using TreeAnnotator 10.5.0 (Bouckaert et al., 2014) with a burn-in of 10%. The BI trees were then used for GMYC species delimitation using the package ‘splits’ (Species Limits by Threshold Statistics) (Ezard et al., 2009) in R 4.4.3 (R Core Team, 2023).

### 2.3 Geometric morphometrics

#### Sampling

For morphometrics, 13 *L. gastrophysus* specimens were analysed (8 collected in the United States and 5 collected in Brazil). North American specimens (corresponding to the voucher numbers MCZ 159678, MCZ 159679, MCZ 159680, MCZ 159681, MCZ 159682, MCZ 159683, MCZ 166186, and MCZ 166187; Table 1) belong to the Ichthyology collection of the Museum of Comparative Zoology (Harvard), in which the digital photos were taken. The five South American specimens (corresponding to samples AD02-AD06; Table 1) were freshly caught from Rio de Janeiro (Brazil) and photographed upon collection.

#### Sample size considerations

The morphometric analyses were based on a relatively small number of specimens (n = 13), particularly for the South American population (n = 5). This reflects constraints that have historically limited morphological studies of the species: *L. gastrophysus* is a demersal species of the continental shelf and slope whose targeted capture in Brazil has been legally regulated since 2005 (Valentim et al., 2007), and specimens suitable for cephalic landmarking are scarce in scientific collections — Caruso (1983) himself cautioned that inferences about the southern population were limited by inadequate sampling. We adopted strict methodological safeguards to mitigate the risks associated with high variable-to-specimen ratios when sample sizes are small. First, discriminant procedures were confined to a low-dimensional space: the Linear Discriminant Analysis (LDA) was applied only to a restricted subset of four principal components, ensuring the number of predictors remained strictly below the size of the smallest geographic group (n = 5), thus avoiding spurious group separation (Bookstein, 2019). Second, to prevent inflated classification rates and resubstitution bias, the accuracy of the discriminant model was evaluated exclusively via leave-one-out (jackknife) cross-validation. Third, supervised morphometric classifications were cross-checked against an independent, unsupervised procedure (Gaussian finite mixture modeling via *mclust*). The conclusions rely on concordance between morphological and molecular evidence rather than on morphometric results alone, as species identification in geometric morphometrics is sensitive to sampling error (Cardini et al., 2021); the accuracy of the landmarking protocol was validated by comparing manual and *in silico* measurements (Section 3.2).

#### Imaging and landmarking

Photographs were taken using a Nikon D5300 equipped with 18-140 mm lenses, incorporating a scale for subsequent size standardisation. Specimens were positioned ventral side down. Landmarking was performed using TPSDig 2.3.1 (Rohlf, 2015) with the 34 2D landmarks previously adopted by Duarte et al. (2004) based on groups of cephalic spines characteristic of *Lophius* species (Figure S1, Supporting Information 2). The landmark corresponding to the end of the vertebral column was excluded from the analysis due to substantial variability observed in the orientation of the posterior margin of the caudal fin in *L. gastrophysus*. Distances between landmarks were also measured manually in three samples to validate the use of measurements performed from photographs. Landmark coordinates were superimposed using a Generalized Procrustes Analysis (GPA) implemented in the *geomorph* package (Adams & Otárola-Castillo, 2013) in R (R Core Team, 2023) to eliminate non-shape biases arising from differences in photo orientation, position, or size.

#### Statistical analyses

All statistical analyses were performed in the R programming environment (R Core Team, 2023). To explore the major axes of shape variation without *a priori* grouping assumptions, a Principal Component Analysis (PCA) was conducted directly on the aligned Procrustes coordinates using the *geomorph* package (Adams & Otárola-Castillo, 2013). To investigate structural variation linked to the population of origin (North America vs. South America), we performed a distance-based Procrustes Analysis of Variance (PERMANOVA) utilizing the RRPP package (Collyer & Adams, 2018), running 10,000 permutations with residual randomization to calculate both global variance explanation and pairwise statistical significance.

A Linear Discriminant Analysis (LDA) was subsequently conducted using the MASS package (Venables & Ripley, 2002) to assess the morphological separation between the two lineages. To prevent artefactual inflation of classification accuracy and model overfitting, the dimensionality of the dataset was reduced prior to the LDA; the discriminant function was computed using a restricted subset of principal components, ensuring the number of predictor variables remained strictly below the sample size of the smallest geographic group. The robustness of the LDA model was validated through a leave-one-out (jackknife) cross-validation procedure, providing an unbiased estimate of classification accuracy and error rates per population. Additionally, an unsupervised clustering analysis using Gaussian finite mixture modeling was performed via the *mclust* package (Scrucca et al., 2016). By treating the restricted principal components as continuous variables without assigning prior geographic labels, this step served as an objective corroboration of the natural grouping structure.

### 2.4 Ethical Statement

This study relied exclusively on preserved museum specimens and tissue samples obtained from dead individuals sourced from commercial fisheries. No live animals were collected, handled, or subjected to experimental procedures during this research. Therefore, specific approval from an institutional animal care and use committee was not required.

## 3 RESULTS

### 3.1 Molecular analyses

#### Molecular divergence and phylogenetic relationships among Lophius species

A total of 243 cytochrome oxidase 1 (*COI*), cytochrome b (*Cytb*), and nuclear rhodopsin (*Rod*) sequences, 84 of which were generated in this study, were used to build maximum likelihood trees (Tables S1-S3, Supporting Information 1).

In the *COI* (Figure 2) and *Cytb* (Figure 3) trees, all seven currently accepted *Lophius* nominal species were recovered as distinct monophyletic clusters, supported by ultrafast bootstrap support > 91%. In addition, both trees recovered Northern and Southern American *L. gastrophysus* as distinct clades (UFBoot support, 100%). The nuclear rhodopsin topology (Figure S2, Supporting Information 2) yielded overall low bootstrap support values, clustering the African *L. vaillanti* with the European species *L. piscatorius* and *L. budegassa*, and the Northern American *L. gastrophysus* with the Southern American *L. gastrophysus*.

**Figure 2.**
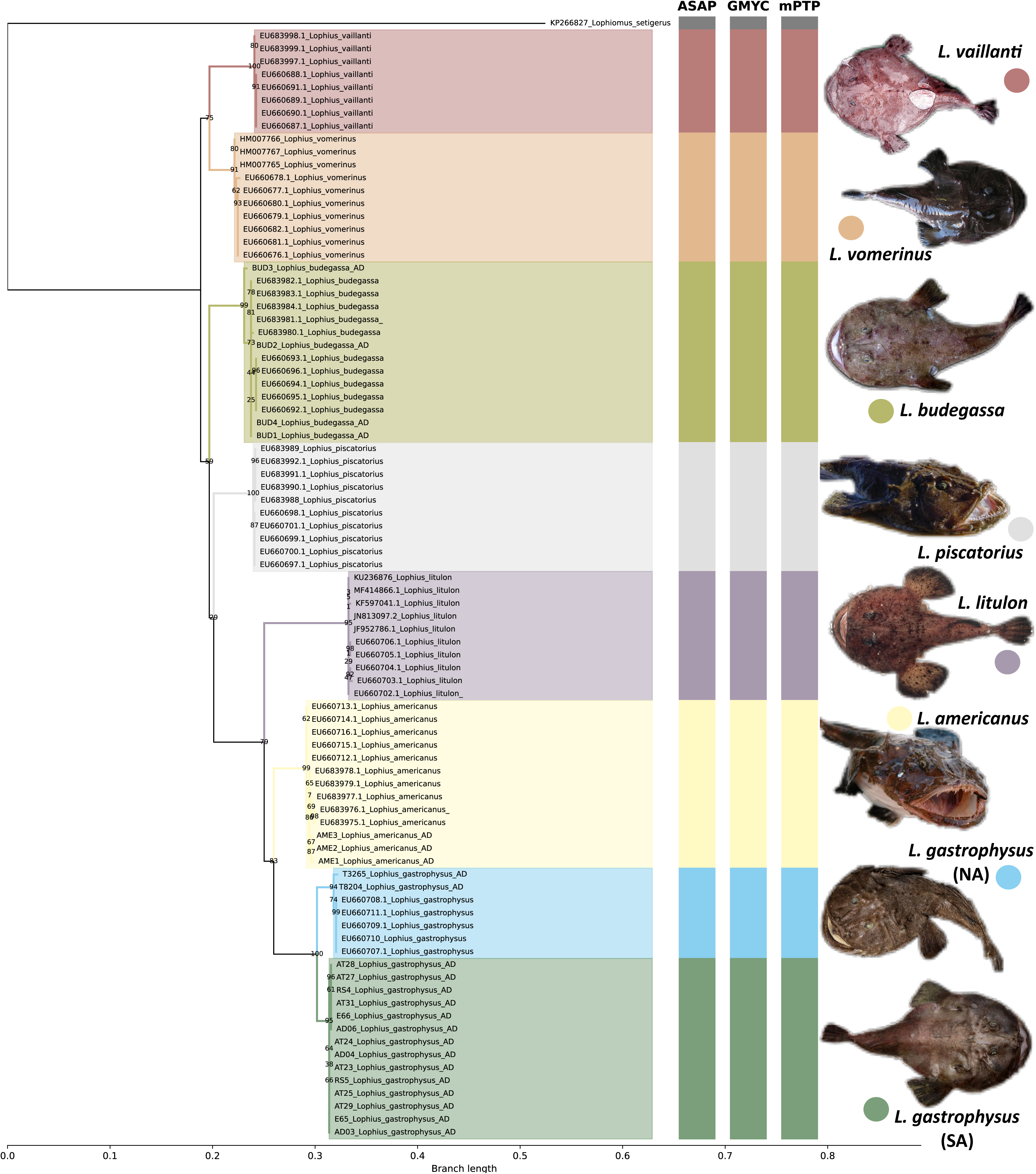
Maximum likelihood phylogenetic tree inferred from Cytochrome c oxidase I (*COI*) sequences. Bootstrap values are indicated as branch labels. Bars indicate the operational taxonomic units (OTUs) recognised by the species delimitation methods. The clade comprising *Lophius gastrophysus* individuals from North American (NA) sampling locations is highlighted in blue, and the clade comprising individuals from South America (SA) is highlighted in green.

**Figure 3.**
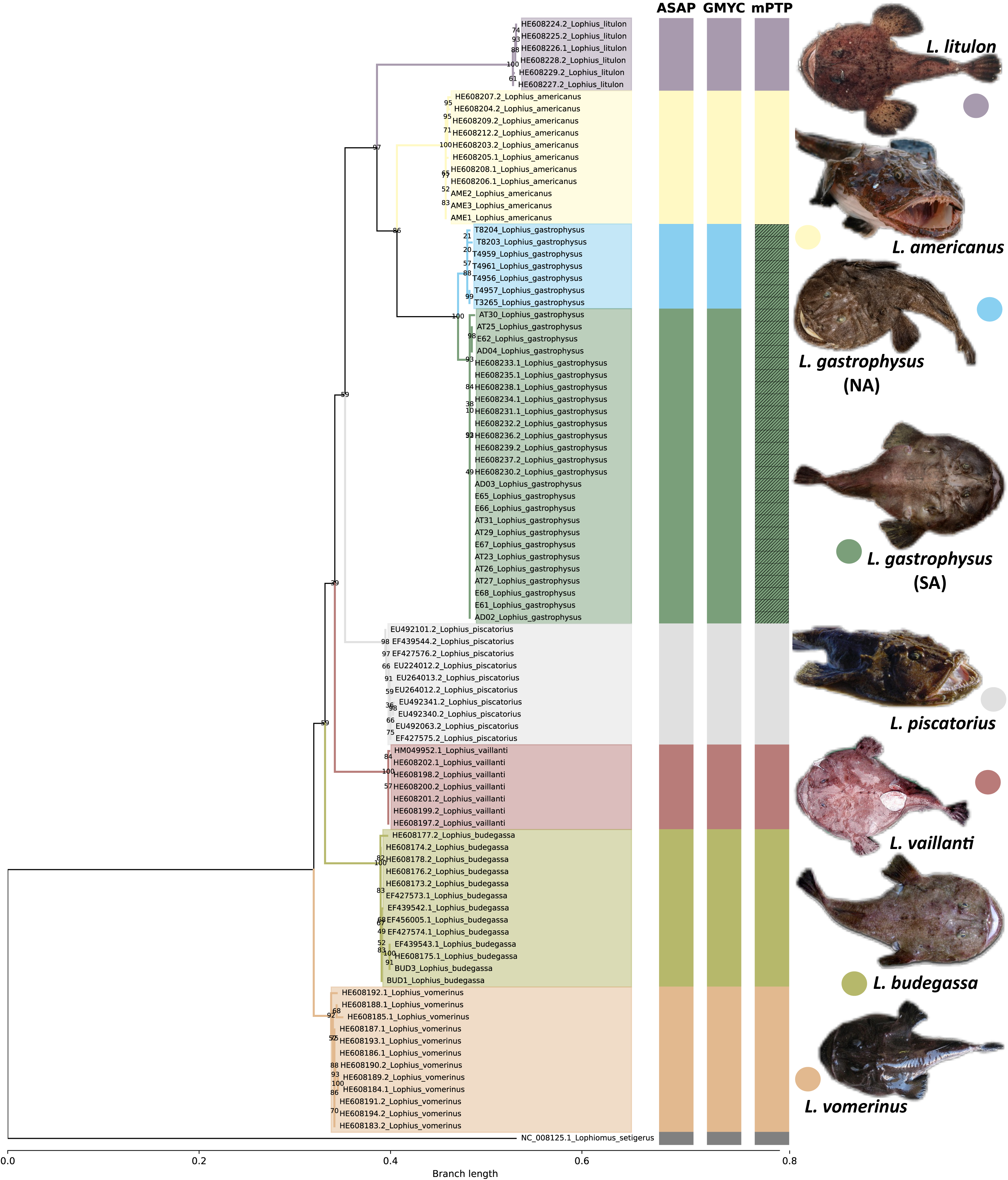
Cytochrome b (*Cytb*)maximum likelihood phylogenetic tree. Bootstrap values are indicated as branch labels. The bars indicate operational taxonomic units (OTUs) recognised by the species delimitation methods. The clade comprising *Lophius gastrophysus* from North American (NA) sampling locations is highlighted in blue, and the South American (SA) clade in green.

The Bayesian phylogenetic trees inferred from *COI* and *Cytb* (Figure S3a and b) sequences presented similar phylogenetic relationships among species, recovering all nominal species as monophyletic and *L. gastrophysus* as two clades (NA and SA). Both trees also revealed the three major biogeographical clades (Euro-African, Asiatic, and American) defined by Grant and Leslie (1993).

The mean intraspecific distance within *Lophius* species for both the cytochrome c oxidase I and the cytochrome b gene was 0.003 (Table S4, Supporting Information 1). Contrastingly, the distance between the North (NA) and South Atlantic (SA) clades was seven times larger (*COI*: p=0.023; *Cytb*: p=0.022; Table S5, Supporting Information 1). The level of divergence observed between NA and SA *L. gastrophysus* was roughly twice the mean intraspecific *COI* distance reported within *Lophiomus setigerus* (<1%) but about half of the *COI* divergence (4.9%) between *Lophiomus setigerus* and its sibling species *Lophiomus immaculioralis* (Chen et al. 2024), indicating that the NA and SA lineages of *L. gastrophysus* have attained a level of mitochondrial differentiation intermediate between intraspecific and interspecific variation in the sister genus.

#### Molecular species delimitation

The three models used to infer species boundaries (ASAP, mPTP and GMYC) showed overall consistent results when applied to the mitochondrial markers (*COI* and *Cytb*), recovering the same eight operational taxonomic units (OTUs): the nominal species *L. piscatorius*, *L. budegassa*, *L. vaillanti*, *L. vomerinus*, *L. litulon*, *L. americanus,* plus the Northern and Southern Atlantic *L. gastrophysus* as two distinct OTUs (Figures 2 and 3).

The ASAP species delimitation analysis based on *COI* sequences identified seven to eight OTUs among *Lophius* sequences across the two partitions with the best ASAP-scores (2.00 and 3.00). For *Cytb*, eight *Lophius* OTUs were consistently identified in the two best-scoring partitions (asap-scores of 1.00 and 4.50).

The GMYC approach also recovered eight OTUs for *COI* (confidence interval 8-9 species, maximum likelihood of null model = 727.0983, maximum likelihood GMYC model = 738.5562, threshold time = -0.0069, p = 1.05 x 10^-5^) and *Cytb* (confidence interval 7-8, maximum likelihood null model = 774.0025, maximum likelihood GMYC model = 786.9008, threshold time = -0.007, p = 2.5 x 10^-6^). Among these eight units, *L. cf. gastrophysus* was split into two distinct taxa (NA and SA), while the remaining OTUs corresponded exactly to the other nominal species.

The multi-rate Poisson Tree Processes (mPTP) delimitation analysis recovered 8 operational taxonomic units (OTUs) from the *COI* dataset. In this analysis, *L. gastrophysus* was subdivided into two distinct OTUs, corresponding to Southern and Northern Atlantic individuals. In contrast, all remaining nominal species (*L. americanus, L. litulon, L. piscatorius, L. budegassa, L. vomerinus,* and *L. vaillanti*) were each recovered as well-defined OTUs. In contrast, the mPTP analysis based on *Cytb* sequences failed to distinguish Northern and Southern Atlantic populations of *L. gastrophysus*. As observed for *COI*, all other nominal species were recovered as independent OTUs in the *Cytb*-based mPTP analysis.

### 3.2 Geometric morphometry

Manual and digital landmarking showed a high degree of correlation (R2 > 96%), and no significant difference was found between manual and digital measurements (p-value = 0.66) (Figure S4, Supporting Information 2).

Principal Component Analysis (PCA) revealed that the first two principal components accounted for 51.91% of the overall shape variance across the dataset, with PC1 and PC2 explaining 31.82% and 20.09% of the variation, respectively (Figure 4a). To test the significance of this morphological differentiation, a distance-based Procrustes Analysis of Variance (PERMANOVA) was conducted. The analysis confirmed a statistically significant structural separation between the Southern (BR) and Northern (EUA) Atlantic L. gastrophysus populations (F = 3.52, p = 0.0023). Geographic origin was a significant driver of shape divergence, accounting for 24.26% of the total morphological variance observed among the specimens. Pairwise comparisons further corroborated this significant separation between the two lineages (d = 0.107, p = 0.0022). To rigorously evaluate the classification of individuals to their respective geographic populations, a Linear Discriminant Analysis (LDA) was employed (Figure 4b). To prevent artefactual inflation of classification accuracy and address the high variable-to-specimen ratio (p > n), the predictive model was strictly limited to four principal components, ensuring the predictor count remained below the smallest group size (n = 5). To validate the phenotypic distinction and avoid the circularity of resubstitution, we implemented leave-one-out (jackknife) cross-validation. The cross-validated LDA model achieved a high global classification accuracy of 92.31%, with an overall error rate of 7.69%. When evaluating the per-group performance, the discriminant function demonstrated robust separation: 100% of the Brazilian (BR) specimens and 87.5% of the United States (EUA) specimens were correctly classified to their true geographic origins.

**Figure 4.**
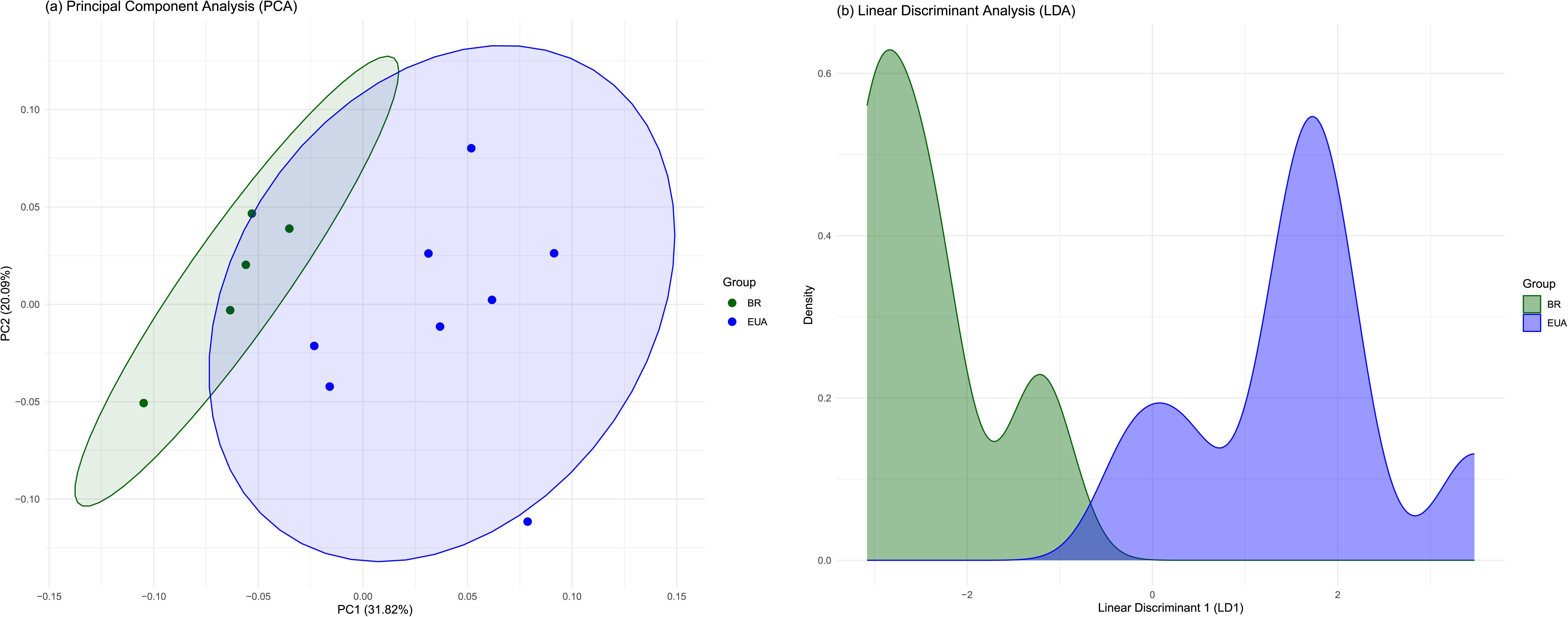
Geometric morphometrics and shape variation of *Lophius gastrophysus* lineages. (a) Principal Component Analysis (PCA) scatterplot showing the distribution of individuals from the Northern Atlantic (EUA, blue) and Southern Atlantic (BR, green) along the first two principal components, which together account for 51.91% of the total shape variance. (b) Density plot derived from the cross-validated Linear Discriminant Analysis (LDA), illustrating the structural separation between the two geographic populations along the first discriminant axis (LD1).

An unsupervised Gaussian finite mixture modelling analysis (mclust) was performed to evaluate the clustering structure without a priori geographic labels. Unlike the LDA, the optimal model (VVI, BIC = 166.37) identified three morphological clusters rather than two strict geographical groups. Individuals from both populations were distributed across these clusters (e.g., Cluster 1 contained 2 BR and 4 EUA specimens). This indicates that while the populations possess distinct, statistically significant morphological signatures capable of being discriminated via directed predictive modelling (LDA), their overall shape variation exhibits natural overlap under completely unsupervised conditions.

## 4 DISCUSSION

The main result of this study is that *Lophius gastrophysus* from the Northern and Southern Atlantic represent phylogenetic lineages that may correspond to distinct species. This conclusion is supported by three complementary lines of evidence: distinct clades recovered in the *COI* and *Cytb* phylogenetic reconstructions, the recognition of Northern and Southern Atlantic individuals as separate Operational Taxonomic Units by automatic species delimitation methods, and morphometric differentiation. Together, these results indicate that *L. gastrophysus*, as currently recognised, comprises at least two independently evolving lineages.

### Genetic distances and phylogenetic inferences

The level of intraspecific variation within *L. gastrophysus sensu lato* was notably high in the mitochondrial genes. At the *COI* gene, the distance (1.14%) was approximately 5 times greater than the average intraspecific distance observed across the genus (0.21%). For the *Cytb* gene, variation within *L. gastrophysus s.l.* (0.8%) was about three times the genus average intraspecific distance (0.3%). The intraspecific variation obtained here for *L. gastrophysus s.l.* is substantially higher than that reported by Espiñeira et al. (2008) based on *COI* sequences, likely due to differences in sampling coverage, as their study included samples exclusively from Florida (USA).

In the sister genus *Lophiomus*, *COI* distances between the sibling species *L. setigerus* and *L. immaculioralis* reach 4.9% (Chen et al., 2024), while mean intraspecific distances within *L. setigerus* remain below 1% (Chen et al., 2024). With 2.3% for *COI* and 2.2% for *Cytb*, the NA–SA clades fall closer to the interspecific end of that range, and both values exceed the 0– 1.83% interval of intraspecific *COI* variation reported for species in a regional barcoding survey (Xing et al., 2018). Model choice (K2P in the *Lophiomus* study versus TN93 here) should not strongly influence these comparisons, as they converge at such low divergences. The figures sit slightly below the 3.4–12.4% window reported for marine sister species on *COI* and *Cytb* (Ramírez-Amaro et al., 2021), which indicates a comparatively recent split. What matters for species delimitation is the internal structure of the pattern: two deeply divergent clades, with within-clade variation several times lower than the between-clade distance, on two independent mitochondrial markers. This combination has repeatedly been the first indication of overlooked species in marine fishes (Durand & Borsa, 2015), and in *L. gastrophysus* it aligns with the delimitation and morphological results in supporting the recognition of the NA and SA lineages as distinct species or, more conservatively, distinct evolutionary lineages.

The mitochondrial maximum-likelihood topologies recovered *L. gastrophysus s.l.* sequences from the Northern and Southern Atlantic populations as two highly diverged and strongly supported (UFBoot support, 100%) monophyletic groups. In contrast, the maximum likelihood tree based on *Rod* sequences did not recover the same groups, which can be mainly attributed to the slower evolutionary rate often observed in nuclear markers (Brown et al. 1979). The *Rod* topology also did not distinguish *L. piscatorius* and *L. vaillanti* as distinct species, underscoring the primary issue of high nucleotide sequence conservation in nuclear regions.

Both *COI* and *Cytb* maximum likelihood topologies for the genus conflict with the mitochondrial phylogeny developed by Espiñeira et al. (2008), which revealed three major clades corresponding to the primary geographical distribution areas occupied by *Lophius* species. Rather than supporting a cohesive Euro-African clade, the *COI* topology obtained here indicates that the European and African species *L. vomerinus* and *L. vaillanti* represent the earliest divergences within the genus, followed by the remaining two species traditionally assigned to the Euro-African clade. In this topology, the American species and the Asian *L. litulon* were recovered as sister groups. The *Cytb* phylogenetic inference yielded a similar departure from the Espiñeira et al. (2008) hypothesis, recovering *L. vomerinus* as the first diversification event within *Lophius*. Together, these results suggest that mitochondrial markers do not uniformly support a biogeographically structured Euro-African clade and point to a more complex evolutionary history within the genus.

Our findings partially align with the biogeographical diversification hypothesis proposed by Grant and Leslie (1993), who, based on allozyme data, identified the African *L. vaillanti* as the first divergence within the genus *Lophius*.

### Molecular species delimitation

The three approaches applied to the mitochondrial datasets (ASAP, GMYC, and mPTP) yielded largely concordant results. Among all nominal extant species of the genus, *L. gastrophysus* was the only one for which mitochondrial data repeatedly supported a taxonomic split. Most delimitation methods recovered the Northern and Southern Atlantic lineages as separate Operational Taxonomic Units (OTUs). The only exception was the multi-rate PTP (mPTP) approach using *Cytb* data, which identified *L. gastrophysus* as a single evolutionary unit. This discrepancy is attributable to the distinct algorithmic framework of the mPTP model, which is inherently more conservative. Designed to mitigate the over-splitting tendencies commonly observed in other tree-based methods (e.g., ASAP and GMYC), mPTP incorporates varying intraspecific coalescent rates (Kapli et al., 2017). Consequently, it often fails to delimit recently diverged lineages exhibiting shallow genetic distances, a limitation particularly pronounced in single-locus mitochondrial datasets like *Cytb* where substitution rates may not have accumulated sufficiently to cross the algorithm’s speciation threshold. Simulation-based analyses corroborate this, indicating that PTP-based methods tend to underestimate species boundaries under conditions of shallow divergence (Luo et al., 2018). Given the robust consensus across all other molecular delimitation methods, combined with the significant morphological differentiation observed, the mPTP result likely represents a conservative underestimation of diversity (false negative) typical of recent divergence events, rather than a true reflection of panmixia.

The delimitation methods can be sensitive to multiple factors, such as incomplete lineage sorting, introgression, or recent hybridisation (Kekkonen & Hebert, 2014). Talavera et al. (2013) showed that area-sampling imbalances across taxa can lead to elevated intraspecific divergence, potentially altering tree structure and increasing the risk of over-splitting in GMYC delimitation. However, it was also noted that sampling intermediate haplotypes did not substantially affect delimitation (Talavera et al., 2013); in that case, the most suitable sampling strategy would primarily include the most extreme haplotypes. Therefore, although one potential limitation of our study is the absence of samples collected from intermediate locations in Central America or Northern Brazil, this specific constraint is likely to have only a limited influence on the GMYC delimitation results.

Because each delimitation method explores only a subset of parameters within the broad parameter space relevant to species delimitation and makes several simplifications, incongruence between the biological units recovered by each method is expected due to the limitations inherent to each analysis (Carstens et al., 2013). Thus, researchers should incorporate data from diverse methods and seek congruent results across them to strengthen confidence in the findings (Carstens et al., 2013). Despite the shortcomings of the three methods applied here, the concordant findings suggest the robustness of the identification of individuals of *L. gastrophysus* from the Northern and Southern Atlantic as distinct species.

The COI distance between the Northern and Southern Atlantic clades (p = 0.023) falls within the range sometimes designated a "grey zone" under universal barcoding thresholds, but this does not weaken the two-lineage hypothesis. COI divergence between congeneric species is strongly taxon-specific (Wiemers & Fiedler, 2007), and fixed thresholds generate both false positives and false negatives (Meyer & Paulay, 2005); the observed distance is nonetheless seven times the mean intraspecific distance across the genus (0.003), and the distance-based delimitation methods used here infer boundaries from distance distributions rather than arbitrary cut-offs (Puillandre et al., 2012). Both mitochondrial genes independently recovered the North–South partition as two fully supported monophyletic clades (UFBoot = 100%), congruent association of independent markers within individuals being the primary molecular criterion for cryptic species delimitation (Padial et al., 2010). The morphometric differentiation of the same populations (PERMANOVA: F = 3.52, p = 0.002; cross-validated LDA accuracy = 92.3%) is an independent line of evidence: every discipline carries a considerable failure rate, and confidence is gained by combining complementary approaches (Schlick-Steiner et al., 2009) and evaluating congruence among independent datasets (Duran et al., 2024). The failure of the nuclear rhodopsin marker to recover the split reflects low resolution at this timescale rather than admixture: rhodopsin did not separate the long-established *L. piscatorius* and *L. vaillanti*, consistent with slower nuclear substitution rates (Martin, 1999) and with the expectation that recently diverged lineages rarely achieve reciprocal monophyly at single loci owing to incomplete lineage sorting (Zhou et al., 2017). Although deep mitochondrial splits do not always imply significant population divergence, and cytonuclear discordance can mislead single-locus delimitation (Dufresnes et al., 2020), concordance between two mitochondrial genes, multiple delimitation methods, and independent morphological evidence indicates that the two lineages are not such an artefact. Genome-wide nuclear data remain the definitive test (Dupuis et al., 2012) and a priority for future work.

### Geometric morphometry

Combining molecular and morphological data often produces more robust results in species delimitation and taxonomy (Will et al., 2005). In line with this, we performed a geometric morphometric analysis using landmarks corresponding to the positions of cephalic spines, as described by Duarte et al. (2004). Conducting morphometric analyses proved crucial, particularly in closely related species with short divergence times, which often do not exhibit significant differences in shape, size, body architecture, or behaviour (Leslie & Grant, 1994; Richman & Price, 1992).

Morphological analysis indicated that the structural arrangement of the head, particularly the displacement in the region of the humeral spines, was a major contributing factor to the primary axes of shape variation. Because these axes were statistically significant in discriminating between Northern and Southern Atlantic *L. gastrophysus* individuals, this morphological divergence constitutes a plausible diagnostic trait. This finding was robustly validated by a Linear Discriminant Analysis (LDA) with leave-one-out (jackknife) cross-validation and corroborated by unsupervised clustering, underscoring the structural distinctiveness of the lineages.

Our results align with those of Leslie and Grant (1994), who similarly suggested two distinct lineages in *L. gastrophysus sensu lato* based on meristic and linear morphometric characters. In their principal component analysis, *L. gastrophysus* exhibited the largest ellipsoid, including samples from the Gulf of Mexico and regions south of the Equator, with individuals from these two sampling locations positioned on opposite sides of the ellipsoid (Leslie & Grant, 1994). As Caruso (1983) and Leslie and Grant (1994) hypothesised, the currently defined species could comprise two evolutionary lineages based on a PCA of linear morphometric measurements.

We successfully validated the accuracy of the digital landmarking method used in geometric morphometrics by comparing its measurements with manually measured pairwise distances between landmarks. Our analysis revealed a high correlation (>96%) and found no significant statistical differences (p > 0.05) between digital and manual landmarks across all examined specimens (Figure S4, Supporting Information 2). Thus, we conclude that the digital landmarking approach was highly accurate.

One limitation warrants mention: the morphometric sample is small (n = 13; southern population n = 5), a direct consequence of the scarcity of intact specimens of a heavily exploited species whose directed capture in Brazil has been legally restricted since 2005 (Valentim et al., 2007). Two strict statistical safeguards were implemented to mitigate this limitation and directly address the risk of spurious group separation associated with high variable-to-specimen ratios (Cardini et al., 2021). First, predictive canonical analyses were restricted to a low-dimensional space of four principal components, ensuring the number of predictors remained strictly below the smallest group size to avoid model overfitting. Second, classification accuracy was assessed exclusively through a leave-one-out cross-validation rather than biased resubstitution.

Even under these conservative constraints, the morphological distinction recovered here remains pronounced: geographic origin explained 24.26% of the total shape variance and discriminated the two populations significantly (PERMANOVA: F = 3.52, d.f. = 1, p = 0.002). Furthermore, the cross-validated LDA achieved a high global accuracy of 92.3%, correctly classifying 100% of the Southern Atlantic and 87.5% of the Northern Atlantic individuals, a signal unlikely to reflect mere sampling noise.

Species identification in geometric morphometrics is nevertheless sensitive to sampling error, as Cardini et al. (2021) documented in extensive datasets; we therefore do not rely on the morphometric result alone, but on its convergence with the phylogenetic and species-delimitation analyses. Notably, while Caruso (1983) had reported morphometric differences between northern and southern *L. gastrophysus* (illicium length, snout tendrils), he cautioned that inadequate sampling of the southern population prevented firm conclusions. Leslie & Grant (1994) subsequently suggested two lineages based on meristic and linear morphometric characters; our geometric landmark data add an independent, robust morphological character set to these earlier lines of evidence. Sampling across intermediate localities, ideally combined with genomic data, remains necessary before any formal taxonomic revision.

### Current-based reproductive isolation hypothesis

Our molecular and morphological data corroborate the hypothesis of two evolutionarily independent North and South American lineages of *Lophius gastrophysus*. Their separation may have resulted from the physical barrier imposed by the South Equatorial Current and the physicochemical barrier constituted by the Amazon River sedimentation plume (Martins et al., 2022). This hypothesis is closely associated with the larval dispersion mode of *L. gastrophysus*, characterised by a pelagic larval phase (Valentim et al., 2007). Pelagic transport of larvae through marine currents may act as a corridor for gene flow, especially within each circulation centre, or restrict gene flow when circulation centres are separated by infrequent movement (Scheltema 1986; Palumbi 1994). The amount of gene flow between populations of species that present this dispersion mode can also be affected by changes in direction, temperature, or speed of the currents, which may ultimately contribute to the isolation of populations that were previously connected or to the isolation of individuals from their original populations (White et al., 2010 Tosetto et al., 2022).

The South Equatorial Current (SEC) flows from east to west along the equator toward South America at an average speed of 1-3 km/h and extends to depths up to 50 m (Caínzos et al., 2023). Upon reaching the Brazilian coast, the SEC splits into three warm, oligotrophic, and saline branches: northern, central, and southern. The northern and central branches converge to form the North Brazil Current (NBC), which flows north-westward along the coast toward the Amazon River (Da Silveira et al., 1994). Meanwhile, the southern branch veers southward and feeds the Brazil Current (BC), which transports warm waters toward the southern Brazilian coast (Dossa et al., 2020).

The latitude of the SEC bifurcation varies seasonally, reaching its southernmost position in July, when NBC-driven transport intensifies, and its northernmost position in November, when BC-driven transport increases (Rodrigues et al., 2007). Data on the reproductive cycle of *L. gastrophysus* along the Brazilian coast are limited (Valentim et al., 2007). During this seasonal window, changes in the South Equatorial Current system favour intensified southward transport by the Brazil Current. This circulation regime may reduce the northward dispersal of early life stages originating from Brazilian populations, limiting larval exchange with Caribbean and northern Atlantic regions. Although direct data on larval dispersal are lacking, the temporal overlap between the reproductive period of *L. gastrophysus* in Brazil and prevailing southward oceanic transport could provide a plausible oceanographic mechanism contributing to geographic isolation between Southern and Northern Atlantic populations.

The Amazon River discharge might form an additional reproductive barrier, generating a low-salinity plume extending over the north Brazilian shelf from 1°S to 5°N (Lentz & Limeburner, 1995). In addition to the Orinoco River outflow, this freshwater plume spans 2,300 km of the northwest coast of South America, physically separating Brazilian and Caribbean reef habitats (Araujo et al., 2022). Consequently, the Amazon plume is considered a significant biogeographic barrier that has contributed to diversification between Brazilian and Caribbean populations (Araujo et al., 2022). According to those authors, the Amazon River also played a primary role in the diversification of the Atlantic reef-fish fauna over the past 2.4 million years, as shown by the correlation between temporal patterns of lineage diversification and periods of higher river sedimentation rates and more frequent sea-level fluctuations in the Pleistocene. Adult *Lophius gastrophysus* live below those barriers and may be little affected by the freshwater and sediment from the Amazon plume. However, they are not very vagile, so their dispersal results mainly from larvae living in shallower waters (Valentim et al., 2007).

One limitation of our study is the lack of samples from intermediate localities. The absence of intermediate sites with admixed or transitional populations can affect species delimitation and may lead to over-splitting, especially in cases of isolation by distance (Mason et al., 2020). However, Dupuis et al. (2012) found no relationship between increased geographic sampling and accurate phylogenetic correspondence in species delimitation studies across taxa, regardless of species range size. Despite these contrasting findings, the lack of intermediate sampling remains a key limitation, and future work should incorporate broader geographic coverage to test whether the North–South split persists across the full distribution of *L. gastrophysus,* especially considering recent records that expanded its known range into the south-western Gulf of Mexico (Del Moral-Flores et al., 2024).

### Implications for fisheries management and conservation

Wildlife conservation depends on accurately identifying biologically significant units and their genetic population structure (Oyler-McCance et al., 2022). The concept of “fish stocks” as management units is widely applied in fisheries science as a metric for strategic planning, implementing stock biomass rebuilding programs, and protecting nursery sites and spawning aggregations (Kerr et al., 2010). Insights from genetic data, alongside morphological evidence, have grown over the last few decades to inform management of endangered fish stocks (Hawkins et al., 2016). Here, we provide genetic and morphometric evidence for identifying at least two distinct management units or, possibly, different phylogenetic species. In this case, actual *L. gastrophysus* stocks may have been overestimated. It has been reported that increased fishing pressure *on L. gastrophysus* populations, driven by increased economic value during the early 2000s, led to stock depletion through overexploitation (Valentim et al., 2007). Hence, our findings could point to a more severe scenario of stock depletion.

## 5 CONCLUSION

Our results support the recognition of two distinct lineages – one in the North and another in the South Atlantic – within *Lophius gastrophysus sensu lato*, as suggested by Caruso (1983). Molecular data from mitochondrial markers consistently distinguished Northern and Southern American *L. gastrophysus* as distinct genetic clusters. This result is supported by the morphological analyses, notably the position of humeral cephalic spines.

Further analyses incorporating more individuals, additional intermediate sampling sites, and broader genomic regions are necessary to validate these findings. Should the Northern and Southern Atlantic clades be recognised as two distinct species, upon taxonomic revision, *L. gastrophysus* shall be retained for the South Atlantic monkfish, given that the original description was based on specimens from Rio de Janeiro, Brazil. In contrast, the northern clade shall be assigned to a different species.

## Acknowledgments

We thank Andrew Bentley (Ichthyology Collection Manager at Kansas University) for assisting in the sample loan process and Meaghan Sorce (Ichthyology Curatorial Assistant at the Museum of Comparative Zoology—Harvard University) for taking the photographs employed in morphometrics. This paper is part of the PhD requirements of Leticia Maria Monteiro Soares at the Biodiversity and Evolutionary Biology Graduate Program of the Federal University of Rio de Janeiro, and through her doctoral fellowship, this study was financed in part by the Coordenação de Aperfeiçoamento de Pessoal de Nível Superior - Brasil (CAPES) - Finance Code 001.

## Author Contributions

A. K. P. D.: Conceptualisation, Methodology, Formal analysis, Writing – original draft, Writing – review & editing.

L. M. M. S.: Formal analysis, Data curation, Writing – original draft, Writing – review & editing.

F. H.: Funding acquisition, Methodology, Writing – original draft, Writing – review & editing.

A. M. S. C.: Conceptualisation, Funding acquisition, Writing – original draft, Writing – review & editing.

## Availability of data and materials

All sequences generated in this study are deposited in GenBank under accession numbers: PZ206986–PZ207026 (rhodopsin), PZ208251–PZ208273 (COI), and PZ252756–PZ252783 (Cytb).

