## Supplementary material for "Unravelling the hidden diversity of the monkfish *Lophius gastrophysus* using integrative taxonomy": Supple_Figures

### Supplementary Figures

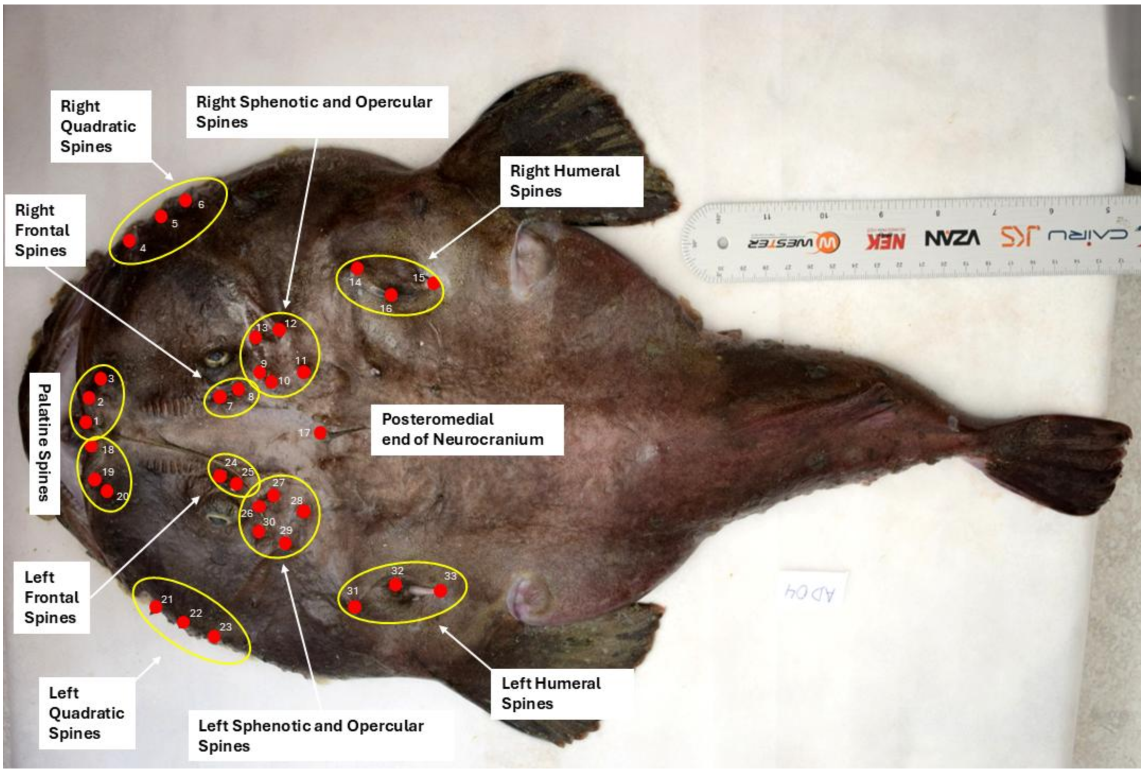


Figure S1. Photograph of the specimen referent to sample AD04 with the representation of the 33 landmarks used in the morphometric analyzes and the corresponding groups of cephalic spines.

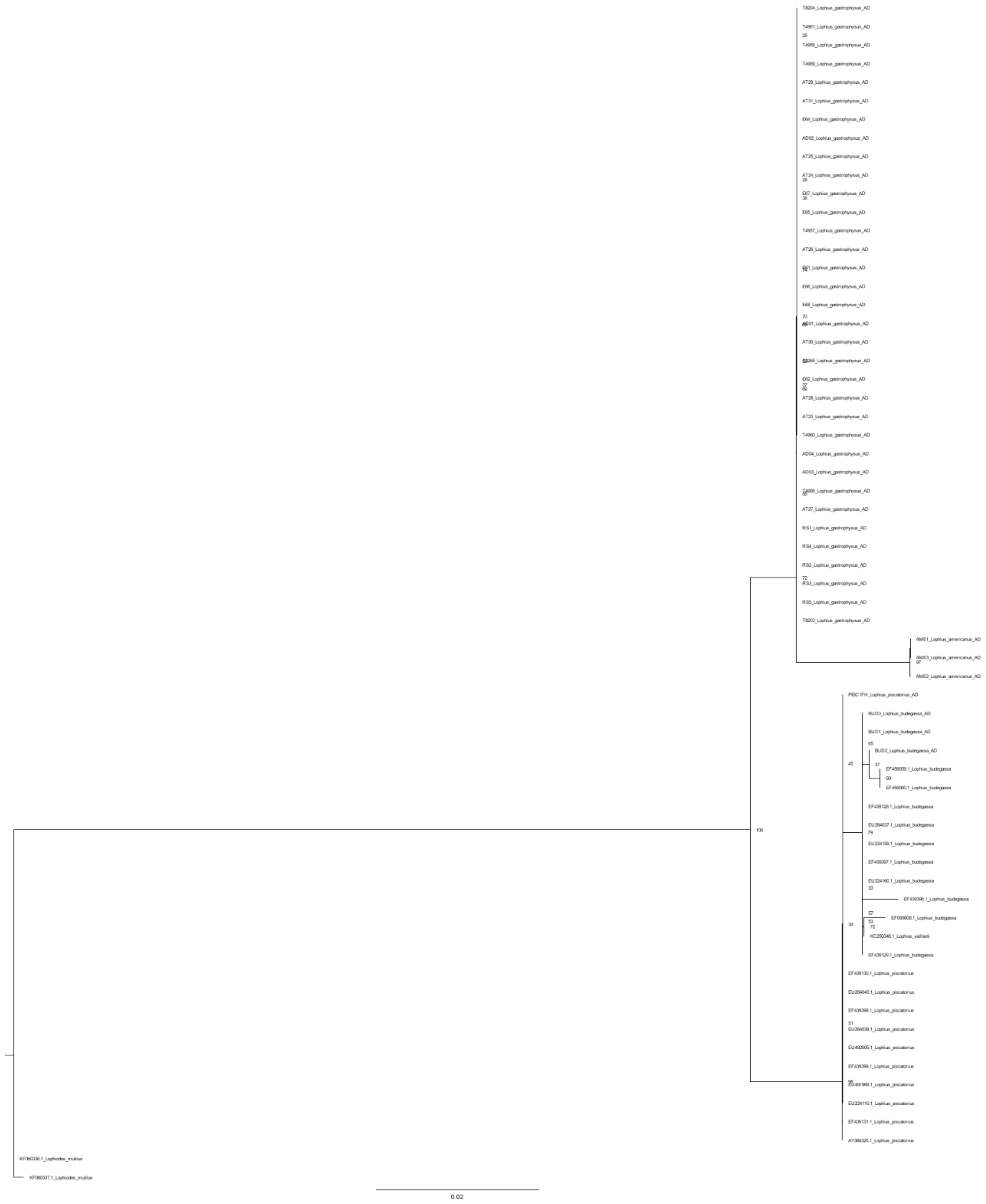


Figure S2. Maximum likelihood Rhodopsin tree. Bootstrap values indicated as branch labels.

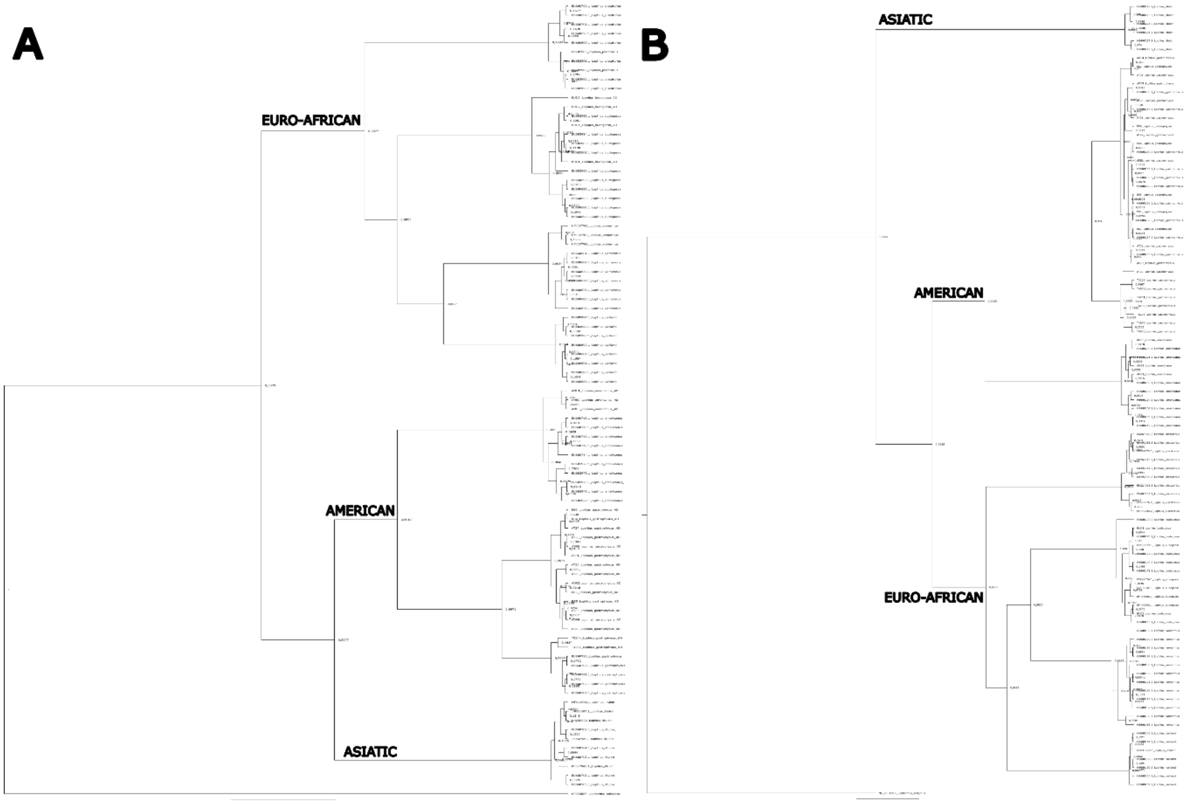


Figure S3. Ultrametric Bayesian trees inferred from *COI* (a) and *cytb* (b) sequences. Bayesian posterior probabilities are indicated at the nodes and biogeographical groups at the branches.

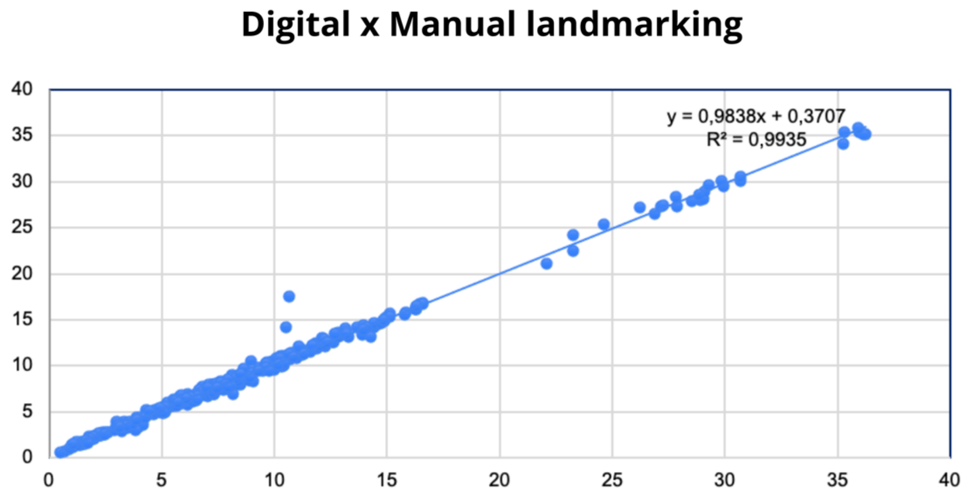


| | |
|---|---|
| Coefficient of determination ($R^2$) | 99.35% |
| Coefficient of correlation (R) | 0.9967 |
| p-value | 0.6655 |

Figure S4. Correlation of distances between landmarks obtained digitally and manually measured for specimens corresponding to samples AD04-AD06 and corresponding p-value.
